# Genetic evidence for a periplasmic protein as a component for a subset of NtrYX two-component systems

**DOI:** 10.1101/2025.11.07.687182

**Authors:** Alexa R. Wolber, Liliana S. McKay, Richard M. Johnson, Zain T. Hameed, Katlyn B. Mote, Steven M. Julio, Peggy A. Cotter

## Abstract

PlrSR, a member of the NtrYX family of two-component regulatory systems (TCSs), is required for the classical bordetellae, including the causative agent of whooping cough, *Bordetella pertussis*, to persist in the lower respiratory tract. The *plrSR* genes are in the middle of a six-gene cluster whose regulation and roles during infection were unknown. *rsmB* and *plrP* are often found 5’ to *plrSR* homologs in β- and γ- proteobacteria, while *trkAH* are often found 3’ to *plrSR* homologs in ⍺-proteobacteria. We investigated these genes to determine if they have a functional link to *plrSR*. We found that this gene cluster does not function as an operon. Rather, it contains two internal promoters: a weaker promoter in the 3’ end of *rsmB* and a stronger promoter in the 3’ end of *plrS.* Additionally, our results indicate that PlrP functions as a third component of the PlrSR TCS. Genetic manipulations of *plrP, plrS,* and *plrR* indicate that PlrP is essential *in vitro* and inhibits PlrS phosphatase activity, likely through PlrS’s PDC domain. Since our results indicate that PlrR can be phosphorylated by another unknown phosphodonor *in vitro*, limiting PlrS phosphatase activity ensures PlrR∼P is not dephosphorylated to lethally low levels. Using natural-host models, we determined that high levels of PlrR∼P are required for *in vivo* survival, and PlrP affects PlrS activity *in vivo*. Given that *plrP* homologs always colocalize with *ntrYX* homologs, we propose that PlrP may fulfill similar functions in other β- and γ-proteobacteria that encode NtrYX- family TCSs, including nonpathogens.

**Importance:** *Bordetella* species, including *B. pertussis*, the causal agent of whooping cough, cause respiratory infections in humans and other animals. Their PlrSR two component regulatory systems, members of the NtrYX family, are required for survival in the lower respiratory tract. We characterized the six-gene cluster that includes *plrS* and *plrR*, identifying one promoter within the first gene that drives expression of the second gene, which we named *plrP*, as well as plrS, and another promoter near the 3’ end of *plrS* that drives expression of *plrR* and the downstream *trkAH* genes. Our data indicate that the *plrP* gene product is an essential third component of the PlrSR TCS, functioning to prevent PlrS from acting as a strong phosphatase *in vitro*. Comparative analyses suggest that PlrP homologs are present, and may function similarly, in NtrYX-family TCSs in other β- and γ-proteobacteria. Our results are important because they provide insight into how bacteria sense and respond to their environment, including those they experience while causing human infection, and this understanding could inform therapeutic and vaccine development.

## Introduction

*Bordetella* species, including the classical bordetellae, are β-proteobacteria that survive in a broad range of environmental and eukaryotic niches. *Bordetella bronchiseptica*, the causative agent of kennel cough in dogs, infects most mammals causing a chronic, often asymptomatic disease. It can also survive long-term in extra-host environments such as filtered pond water(1). By contrast, *B. pertussis* and *B. parapertussis*_hu_, which evolved from a *B. bronchiseptica*-like ancestor, are respiratory pathogens that survive only in the human host (causing whooping cough) and briefly during transmission. Despite high vaccination rates, whooping cough cases are on the rise globally, and better vaccination strategies are needed (2, 3).

All known protein virulence factor-encoding genes in *Bordetella* spp, including those that encode the components of current acellular vaccines, are regulated by the two-component regulatory system (TCS) BvgAS, which has been regarded as the “master virulence regulator” (4–8). BvgAS controls at least three distinct phenotypic modes. Bvg^+^ mode, produced when BvgAS activity is high and virulence-associated genes (*vags*), such as those encoding toxins and adhesins, are maximally expressed, is both necessary and sufficient for bacterial persistence *in vivo* (8–12). Bvg^−^ mode, produced when BvgAS is inactive and virulence-repressed genes (*vrgs*), required for flagella synthesis and motility and other phenotypes, are maximally expressed, does not appear to occur *in vivo* but is required for *B. bronchiseptica* survival in nutrient-limited environments (9, 13, 14). The Bvg^i^ mode is produced when BvgAS is partially active and is hypothesized to occur during transmission between mammalian hosts (15, 16).

In 2011, another TCS, PlrSR, was discovered to be required for *B. bronchiseptica* persistence in the lower respiratory tract (LRT) of rats and mice (17, 18). In addition, BvgAS activity was found to be dependent on PlrSR in the LRT (18), suggesting a functional link between the two TCSs. However, PlrSR is not only required in the LRT because of its effect on BvgAS activity. Even when BvgAS is constitutively active, a mutant that produces non-functional PlrS (Δ*plrS*) fails to survive in the LRT (18).

Therefore, PlrSR must regulate one or more BvgAS-independent genes that is/are required for persistence in the LRT.

PlrS is a histidine sensor kinase (HK) protein with a predicted periplasmic PhoQ-DcuS-CitA (PDC) sensory domain followed by cytoplasmic HAMP, PAS, HK, and HATPase domains. PlrS has a motif (HEIKN) at its primary site of phosphorylation (H521) that is characteristic of sensor kinases that can act as both a kinase and a phosphatase towards their cognate response regulator proteins (19, 20). PlrR is a typical response regulator protein with a N-terminal receiver domain and C-terminal DNA-binding domain. A *B. bronchiseptica* mutant encoding PlrS with glutamine substituted for H521 (H521Q) is as defective as a PlrS mutant missing aa 5-198 (Δ*plrS*) for persistence in the LRT (17), indicating that phosphorylated PlrS, and thus, presumably, phosphorylated PlrR, is required *in vivo*. While the H521Q mutant is defective for kinase activity *in vitro* (21), its phosphatase activity is unknown. The asparagine residue at position H+4 (N525) is required for phosphatase activity, and biochemical analyses indicate that PlrS with alanine substituted for N525 (N525A) may also be partially defective for kinase activity (21).

PlrSR is a member of the NtrYX family of TCSs that are widely distributed in proteobacteria and involved in regulating nitrogen metabolism, cell envelope processes, respiratory enzymes, and iron homeostasis (22–29). *ntrYX* homologs appear to have evolved from *ntrBC*, which encode a TCS important in controlling nitrogen metabolism in proteobacteria (30, 31). NtrYX systems evolved further such that homologs in ⍺- proteobacteria differ from those in β-proteobacteria. Specifically, NtrX response regulator proteins of β-proteobacteria lack the AAA+ domain that is present in NtrX proteins of ⍺-proteobacteria (32). Moreover, the *ntrYX* genes in β-proteobacteria are 3’ to two highly conserved genes: one predicted to encode an RNA methyltransferase (*rsmB*) and one predicted to encode a periplasmic proline-rich domain of unknown function protein (which we are calling *plrP*) (32). The role of this highly conserved genetic organization is currently unknown. Our analyses revealed genetic evidence that *plrP* is involved in PlrSR signaling, and we hypothesize it may be a conserved component of NtrYX TCSs in other β-proteobacteria.

## Results

### *plrSR* are located within a six-gene cluster with operon-like structure

The *plrSR* genes are in the middle of a cluster of genes oriented in the same transcriptional direction (Fig. 1A; BB0262-0267). 5’ to *plrS* are two genes: *rsmB*, predicted to encode an rRNA methyltransferase, and BB0263, here named *plrP*, predicted to encode a protein containing a domain of unknown function. The *rsmB* homolog from *Escherichia coli* encodes a product that methylates C697 of the 16s rRNA (m^5^C967), a residue involved in interactions with tRNA (33–35). By stabilizing the interaction between the pre-initiation complex and the initiating tRNA bearing methionine, m^5^C967 has been shown to impact the efficiency of translation initiation and thus the overall bacterial proteome (36). *plrP* is predicted to encode a 203 amino acid protein containing DUF4390. Based on SignalP analysis, this putative protein contains a signal sequence (amino acids 1-25) for exportation to the periplasm (37).

**Figure 1.**
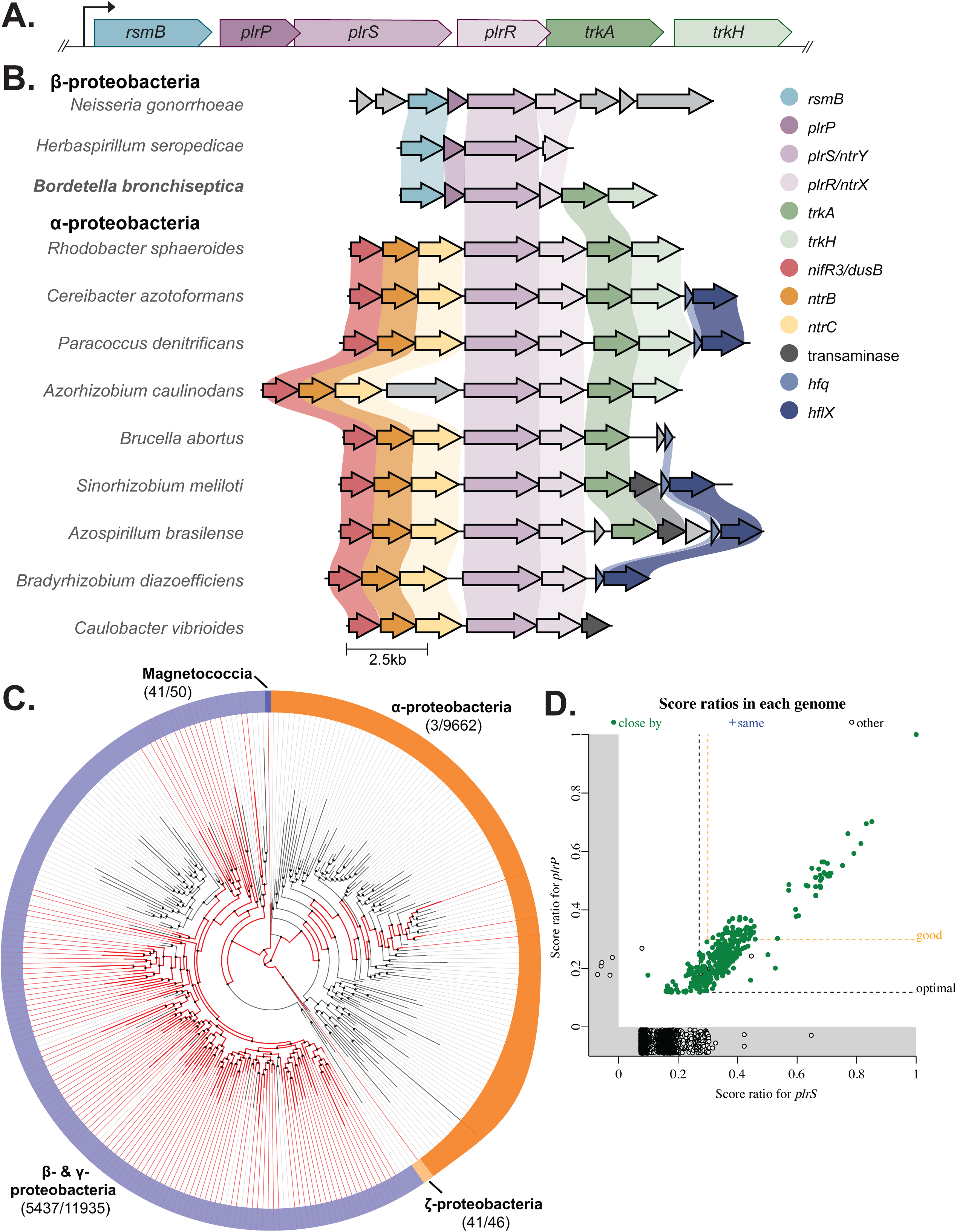
Conservation of genes surrounding *ntrY* and *ntrX* homologs in proteobacteria. (A) Schematic of the *plrSR* gene cluster in *Bordetella* spp containing locus BB0262-BB0267. *plrP* and *plrS* overlap by 4 base pairs, as do *plrR* and *trkA*. (B) Schematic showing the organization of the gene region surrounding *ntrYX* homologs, centered on the *ntrY* homolog. Genes that are predicted to encode the same type of protein are similarly colored. Shaded lines connecting genes indicate >30% sequence identity. (C) Distribution of genes encoding DUF4390-containing proteins within proteobacteria. Red lines represent orders that contain genomes with genes encoding DUF4390-containing proteins. Gray lines represent orders that do not. (D) Gene presence/absence plot of *plrS* homologs vs *plrP* homologs. Score ratios closer to 1 indicate a greater degree of homology to the queried genes. A score ratio below 0 indicates no homolog was detected in that genome. Solid green dots indicate the *plrS* and *plrP* homologs are found within 5kb of each other in the genome. Orange dashed line indicates the arbitrary threshold for good homology (30% of the maximum bit score).

The two genes 3’ to *plrSR* (*trkA* and *trkH*) are predicted to encode a potassium transporter. Studies with TrkA and TrkH from *Vibrio parahymolyticus* indicate that TrkH forms the membrane channel through which potassium ions can be transported, while TrkA remains cytosolic, regulating ion flux through TrkH by changing conformation in response to intracellular ADP/ATP levels (38).

The fact that all six genes in the *plrSR*-containing cluster are oriented in the same transcriptional direction with minimal or no intergenic sequence between predicted start and stop codons suggests they form an operon. The gene 5’ to *rsmB* is oriented in the opposite transcriptional direction with a 107 base pair (bp) intergenic region. While the gene 3’ to *trkH* is in the same transcriptional direction as the gene cluster, 79 bp separate the two genes. There are only 18 bp between the predicted stop codon of *rsmB* and the predicted start codon of *plrP,* and nine bp between the predicted stop codon of *plrS* and the predicted start codon of *plrR*. The 3’ end of *plrP* overlaps with the 5’ end of *plrS* by four bp, and the 3’ end of *plrR* overlaps with the 5’ end of *trkA* by four bp. The largest intergenic region, 48 bp, is between the predicted stop codon of *trkA* and predicted start codon of *trkH*. Based on these features, we hypothesized that there is a promoter 5’ to *rsmB* and that the six genes are expressed as an operon.

### *rsmB, plrP*, and *plrSR* are colocalized in β- and γ-proteobacteria

A previous study examined the genes surrounding *ntrYX* and found a distinction between α- and β-proteobacteria (32). In ⍺-proteobacteria, *ntrYX* was 3’ to *ntrBC,* genes encoding another closely-related TCS. In β-proteobacteria, *ntrYX* was 3’ to an *rsmB* homolog and a gene encoding a proline-rich DUF4390-containing protein like PlrP. To further assess this evolutionary distinction, we compared the gene neighborhood structure of NtrYX-encoding systems within twelve bacterial species, including *B. bronchiseptica,* where the TCS has been studied, using the gene cluster comparison program CAGECAT clicker (39) (Fig. 1B). As reported, the three β-proteobacteria, including *B. bronchiseptica*, had *rsmB* and *plrP* homologs 5’ to their *plrSR* homologs.

The nine ⍺-proteobacteria had *ntrBC*, as well as a *nifR3* or *dusB* homolog, 5’ to their *plrSR* homolog. In both the α- and β-proteobacteria, the genes 3’ to their *plrSR* homologs varied. However, *trkA* and *trkH,* which are 3’ to *plrSR* in *B. bronchiseptica,* were also located 3’ to *plrSR* homologs in seven and four ⍺-proteobacteria species, respectively, but not in other β-proteobacteria.

To assess the relationship between the six genes within the *plrSR* gene cluster in a wider selection of genomes, we examined the rate of co-occurrence of these genes using the *fast.genomics* database (40). First, we examined the neighborhood architecture surrounding the top 200 closest homologs to *plrS*. Out of the 198 with complete sequencing of the region, 195 (98%) had homologs of *rsmB* and *plrP* 5’ to the *plrSR* homologs. By contrast, only 53 (27%) had *trkA* and 46 (23%) had *trkH* 3’ to the *plrSR* homologs, further indicating a weaker conserved relationship between *plrSR* and *trkAH* than between *plrSR, rsmB,* and *plrP*.

Given the conservation of the genes 5’ to *plrSR*, we further examined the distribution of *rsmB* and *plrP* within proteobacteria using AnnoTree, a tool for visualizing the distribution of genes across large phylogenetic trees (41). Genes encoding proteins containing DUF4390 were found within the genomes of β- and γ-proteobacteria, as well as Magnetococcia and ζ-proteobacteria, but not within the genomes of ⍺-proteobacteria (Fig. 1C). Of the β- and γ-proteobacteria included in AnnoTree, a minority (37%) encoded a *plrP* homolog. *rsmB*, however, was widely distributed; 94.3% of all proteobacteria examined encoded at least one copy (Supplemental Fig. 1A).

We then focused our analysis on the co-occurrence of *rsmB, plrP,* and *plrSR* across the entire *fast.genomics* database. *rsmB* homologs were widely distributed even outside of proteobacteria and were found within 76% of the genomes queried, versus the 47% of genomes containing a *plrS* homolog (Supplemental Fig. 1B). While many genomes contained either *rsmB* or *plrS* homologs, the likelihood of encoding both genes increased as the degree of homology increased. Using a cut-off of 30% of the maximum bit score as an indicator of good homology, 336 *plrS* and 490 *rsmB* homologs were identified. 313 of these genomes contained good homologs of both genes and 311 (99.4%) of these homologs were located within 5kb of each other, indicating a conserved functional relationship.

Within the genomes queried, only 446 (6.1%) contained *plrP* homologs. However, 441 of those 446 genomes (98.9%) also contained intact *plrS* homologs (Fig. 1D). The other 5 genomes with *plrP* homologs contained incomplete sequences or pseudogenes with homology to *plrS*. Of the 441 genomes with annotated homologs of both *plrP* and *plrS*, 437 (99.1%) contained both genes within 5kb of each other. Again, using a cut-off of 30% of the maximum bit score as an indicator of good homology, 336 *plrS* and 94 *plrP* homologs were identified. Of the 94 genomes containing good *plrP* homologs, all 94 (100%) also contained a good *plrS* homolog and in every case these homologs were within 5kb of each other. Manual examination of the 336 *plrS* homologs with good homology determined that 333 of these homologs (99.1%) had a gene encoding a DUF4390-containing protein 3’ to it, even if these genes were not determined to have homology to *plrP*. While the vast majority (94%) of these good PlrS homologs were found in the genomes of β-proteobacteria, specifically Burkholderiales, this cut-off also included γ-proteobacteria, which had the same neighborhood architecture. This colocalization of *plrS* and *plrP* homologs strongly implies that *plrP* or the protein it encodes interacts in some capacity with *plrS* or PlrS and that this interaction is important to the function of the TCS in β-proteobacteria.

### RNAseq data suggests internal promoters 5’ to plrP and plrR

Attempts to delete *plrR* or sequences near the 3’ end of *plrS* have been unsuccessful, suggesting that *plrR* is essential *in vitro* and that its expression is driven by a promoter within the 3’ end of *plrS*. To investigate this hypothesis, we analyzed RNAseq data across the *rsmB, plrP, plrS, plrR, trkA, trkH* region from a previous study (42). Among all the conditions assessed (ambient air (∼21% O_2_), 5% O_2_, 2% O_2_, and 20% O_2_ 5% CO_2_), few or no transcripts were detected for *rsmB* (Fig. 2, Supplemental Fig. 2). A moderate number of transcripts was detected at the 5’ end of *plrP*, reaching a peak ∼200 bp within the gene and extending through *plrS* (Fig. 2). Many more transcripts were detected at the 5’ end of *plrR*, continuing, and decreasing, through *trkA* and *trkH* (Fig. 2). These data suggest that, under the conditions tested, *rsmB* is not expressed, that there is a weakly active promoter within the 3’ end of *rsmB*, and that there is a strongly active promoter within the 3’ end of *plrS*.

**Figure 2.**
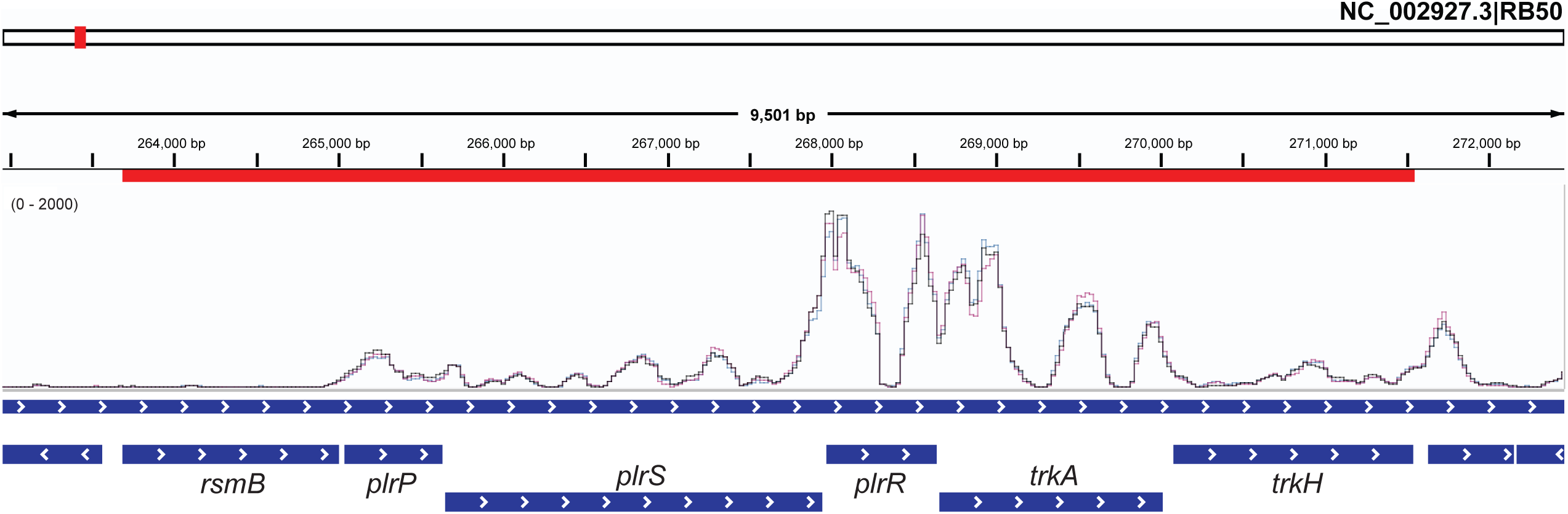
RNAseq analysis of the *plrSR* gene cluster suggests two internal promoters. Graph of RNA transcripts measured using RNAseq from *B. bronchiseptica* samples that were grown in ambient air conditions(42). Data was visualized using the Integrative Genomic Viewer(51).

### P_plrP_ and P_plrR_ are active in vitro

To determine if sequences within the 3’ end of *rsmB* and the 3’ end of *plrS*, as well as those 5’ to *rsmB* (Figure 3), contain promoter activity, we cloned DNA fragments from each region 5’ to the promoterless *gfp* gene in plasmid pMAB*gfp*, delivered the putative promoter-*gfp* cassettes to the chromosomal *att*Tn*7* site in wild-type and Δ*plrS* bacteria, and measured fluorescence after growing the strains in Stainer-Scholte (SS) medium (standard liquid growth medium for *Bordetella* spp) with and without the addition of 40 mM MgSO_4_. (40 mM MgSO_4_ inactivates BvgS (43).) As a control to show that 40 mM MgSO_4_ modulated BvgAS activity in our experiments, we included a strain containing a P*_bvgA_*-*gfp* fusion. *bvgAS* is positively autoregulated, and in addition to BvgAS being inactive in SS medium containing 40 mM MgSO_4_, BvgAS activity is somewhat diminished when PlrS is inactive *in vitro* (18). Consistent with these previously published reports, the wild-type strain containing the P*_bvgA_*-*gfp* fusion was highly fluorescent in SS medium and minimally fluorescent in SS medium containing 40 mM MgSO_4_, and the Δ*plrS* strain containing the P*_bvgA_*-*gfp* fusion was highly fluorescent in SS medium but less so than the wild-type strain, and minimally fluorescent in SS medium containing 40 mM MgSO_4_.

**Figure 3.**
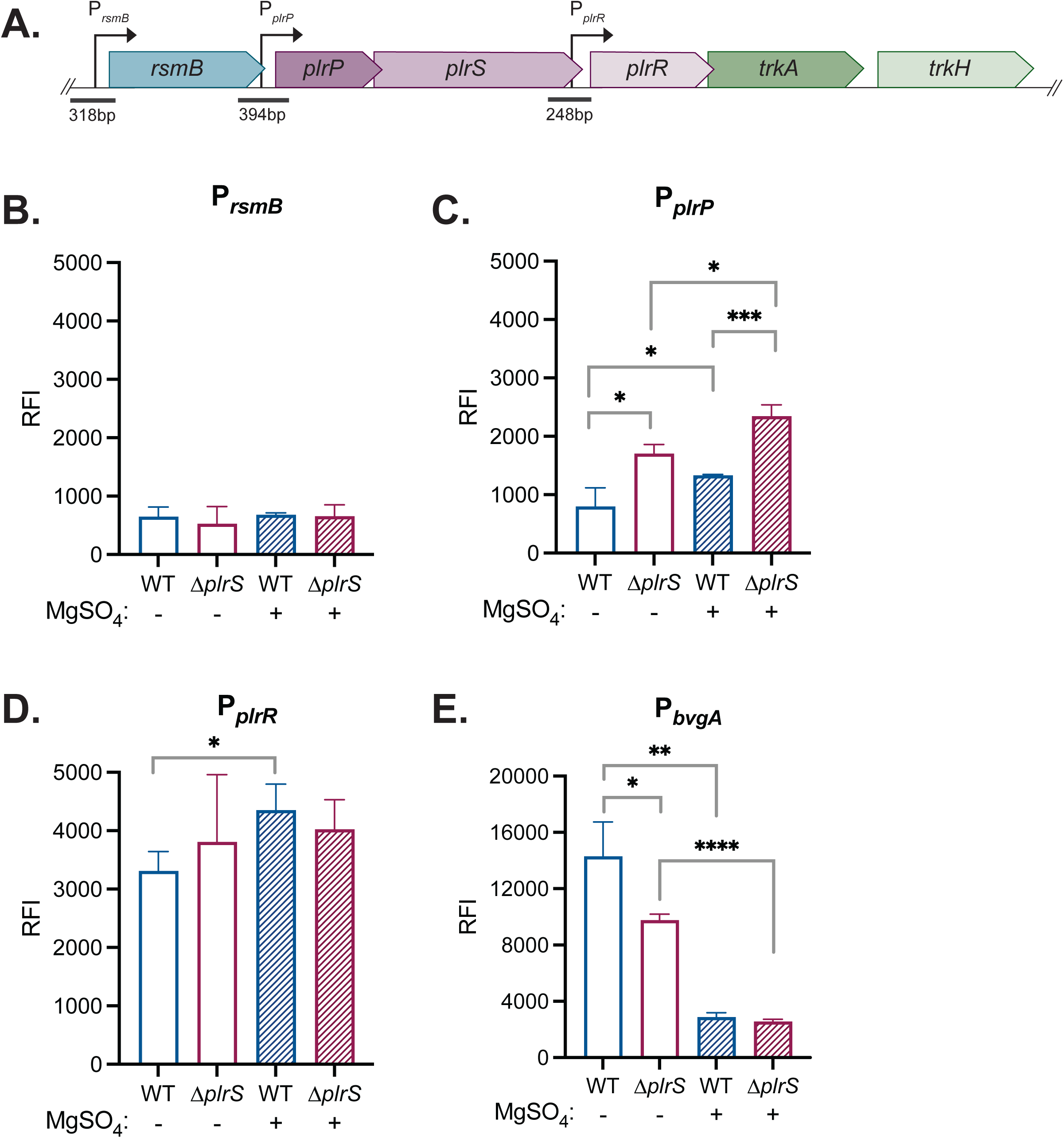
P*_plrP_* and P*_plrR_* are active under standard *in vitro* growth conditions. (A) Schematic of the *plrSR* gene cluster in *Bordetella* spp containing locus BB0262- BB0267. Putative promoter regions are annotated with an arrow indicating the direction of transcription. Fragments cloned to the promoterless *gfp* gene are annotated with a line and the corresponding size in base pairs. Strains containing P*_rsmB_*-*gfp* (B), P*_plrP_*-*gfp* (C), P*_plrR_*-*gfp* (D), P*_bvgA_*-*gfp* (E) cassettes at the chromosomal *att*Tn*7* site in wild-type and Δ*plrS* bacteria were grown in SS medium with and without the addition of 40mM MgSO_4_. GFP fluorescence (excitation: 485nm, emission: 535nm) and OD_600_ were measured after 16-18 hours of growth, and relative fluorescence intensity (RFI) was calculated by dividing GFP fluorescence by OD_600_ values. Statistical significance, as determined using unpaired Student’s t-test, is indicated as *, p< 0.05; **, p<0.01; ***, p<0.001; ****, p<0.0001.

Fluorescence of strains containing the P*_rsmB_*-*gfp* fusion was minimal under all conditions, consistent with the 300 bp 5’ to *rsmB* not containing a promoter that is active in SS medium, with or without the addition of 40 mM MgSO_4_.

Fluorescence of the wild-type strain containing the P*_plrP_*-*gfp* fusion was low in SS medium and about two-fold higher in SS medium containing 40 mM MgSO_4_, suggesting a weak promoter that is more active in the Bvg^−^ mode than the Bvg^+^ mode. The Δ*plrS* strain was moderately fluorescent in SS medium and slightly more fluorescent in SS medium containing 40 mM MgSO_4_. These data indicate that the 400 bp region 5’ to *plrP* contains a promoter that is weakly active under the conditions tested, and that is somewhat negatively regulated by both PlrS and BvgAS. By contrast, the wild-type and Δ*plrS* strains containing the P*_plrR_*-*gfp* fusion were highly fluorescent under all conditions tested. These data indicate the presence of a promoter within the 3’ end of *plrS*.

Overall, these data indicate that *plrP* and *plrS* are transcribed from a promoter within the 3’ end of *rsmB* that is weakly active under the conditions tested, and that *plrR*, *trkA*, and *trkH* are transcribed from a promoter within the 3’ end of *plrS* that is moderately active under the conditions tested.

### *rsmB*, *trkA*, and *trkH* are not required for *B. bronchiseptica* persistence during murine infection

To determine if *rsmB*, *trkA*, and *trkH* contribute to infection, we constructed strains with in-frame deletion mutations in each gene and compared them to wild-type *B. bronchiseptica* for their ability to persist in the murine respiratory tract. Mice were inoculated intranasally with 7.5×10^4^ CFU, and bacterial burdens in the nasal cavity, trachea, and right lung lobes were determined at days 0, 1, and 3 post-inoculation. As previously shown (17, 18), the Δ*plrS* mutant was severely defective for persistence relative to the wild-type strain in the LRT (trachea and lung), but not the nasal cavity (Fig. 4). The Δ*rsmB* and Δ*trkAH* mutants were recovered at levels similar to those of the wild-type strain in the nasal cavity, trachea, and lung at every timepoint (Fig. 4), indicating that neither *rsmB* nor *trkAH* are required for *B. bronchiseptica* persistence during infection in this murine model.

**Figure 4.**
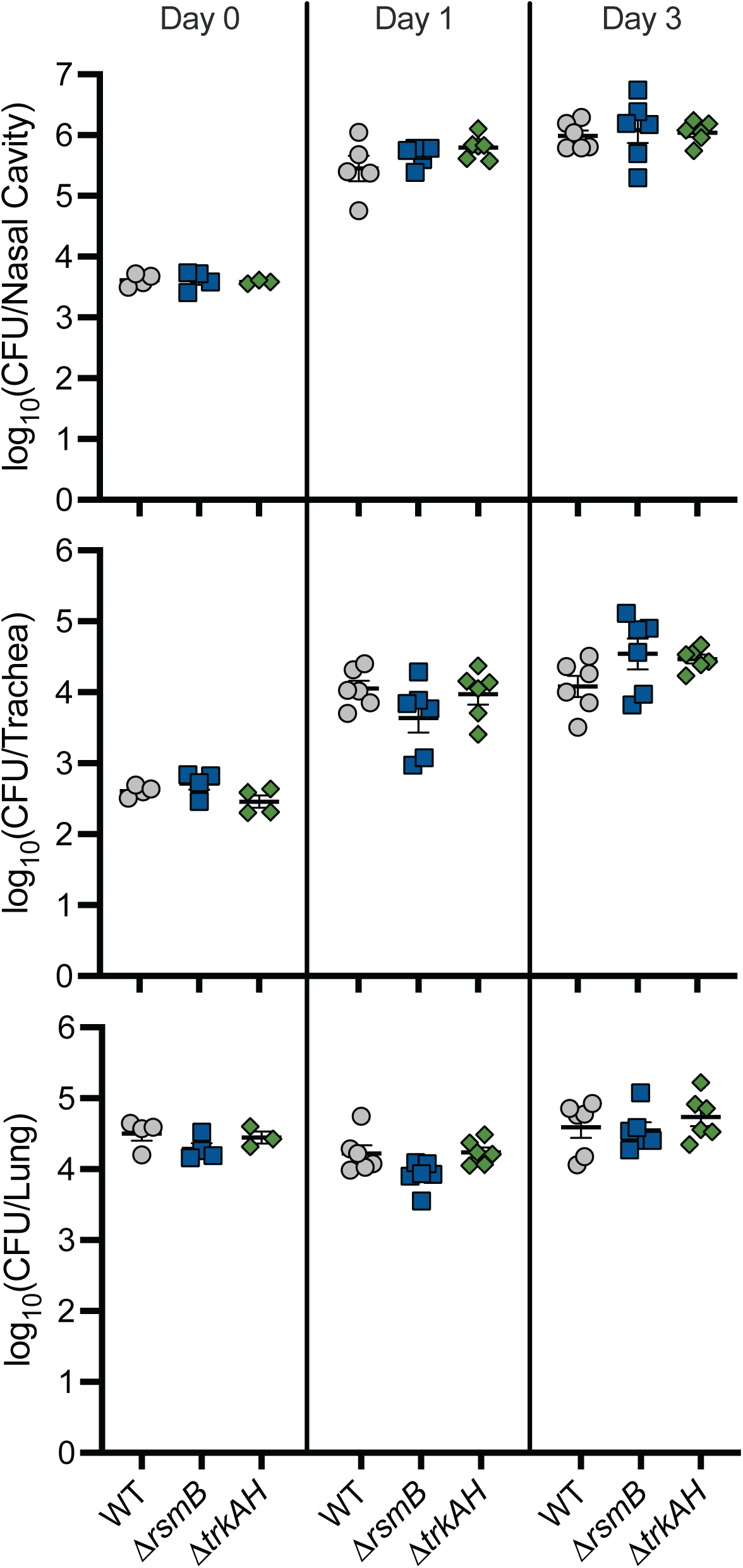
*rsmB*, *trkA*, and *trkH* are not required for *B. bronchiseptica* persistence in the murine respiratory tract. Bacterial burden over time within the nasal cavity (top), trachea (middle), and right lung lobes (bottom) of mice infected with 7.5×10^4^ CFU of wild-type (WT), Δ*rsmB*, or Δ*trkAH* bacteria. n=4 for day 0, and n=6 for days 1 and 3, with each symbol representing a single mouse.

### *plrP* is essential *in vitro* when *plrS* is unmutated

To determine if *plrP* is required for persistence in the LRT, we attempted to construct a strain containing an in-frame deletion in *plrP.* We used an allelic exchange plasmid designed to delete codons 127-600, leaving 12 bp at the 3’ end of *plrP*, which overlaps by four bp with *plrS* (Fig. 1A). We were able to obtain co-integrants with this plasmid, but all colonies (>64 screened) obtained after growth with no antibiotic selection and then plating on agar containing 20% sucrose (i.e., bacteria in which the plasmid has recombined out of the chromosome) contained wild-type *plrP*, suggesting that *plrP* is essential *in vitro*. Using the same allelic exchange plasmid, we were able to delete *plrP* in the PlrS_H521Q_ strain, in which PlrS is unable to be phosphorylated (21), as well as the PlrS_N525A_ strain, in which PlrS is defective for phosphatase activity (21).

Using this allelic exchange plasmid, we could not attempt to delete *plrP* from the Δ*plrS* strain because the Δ*plrS* strain lacks appropriate homologous sequences. We were also able to delete *plrP* in a strain in which *plrR* was deleted from the native site and *plrR* in which the codon for D52 (the primary site of phosphorylation in PlrR) was replaced with a codon for glutamic acid (PlrR_D52E_), a substitution that is predicted to function as a phosphomimetic. Together, these observations suggest that *plrP* is essential *in vitro* in a manner that is dependent on PlrS functionality, and they support the hypothesis that PlrR must be phosphorylated, at least at a low level, *in vitro*, and that PlrP prevents PlrS from fully dephosphorylating PlrR∼P *in vitro* (i.e., that, during growth in SS, PlrP keeps PlrS in ‘kinase mode’).

### *plrP* essentiality requires the PlrS PDC domain

Because PlrP is predicted to contain a signal sequence that directs it to the periplasm, we hypothesized that PlrP affects PlrS activity by interacting with the PlrS PDC domain. We constructed a strain in which codons 463-720 of *plrS* are deleted, resulting in deletion of just the PDC domain. This strain has no obvious growth or colony phenotype *in vitro*, similar to the Δ*plrS* strain and the PlrS_H521Q_ strain. Unlike the case with wild-type bacteria, we were able to delete *plrP* in the ΔPDC strain, indicating that PlrP essentiality *in vitro* requires the PDC domain of PlrS, supporting the hypothesis that PlrP affects PlrS activity via the PlrS PDC domain.

### The PlrS PDC domain is not required for *B. bronchiseptica* persistence in the lower respiratory tract, and if *plrP* plays a role *in vivo*, its role requires the PlrS PDC domain

We compared the ΔPDC and ΔPDC Δ*plrP* strains for their ability to persist in the murine LRT. The mutants were recovered from all sites in the respiratory tract at levels similar to those of wild-type bacteria (Fig. 5), indicating that the PlrS PDC domain is not required *in vivo*, and that if PlrP plays a role during infection, that role requires the PlrS PDC domain.

**Figure 5.**
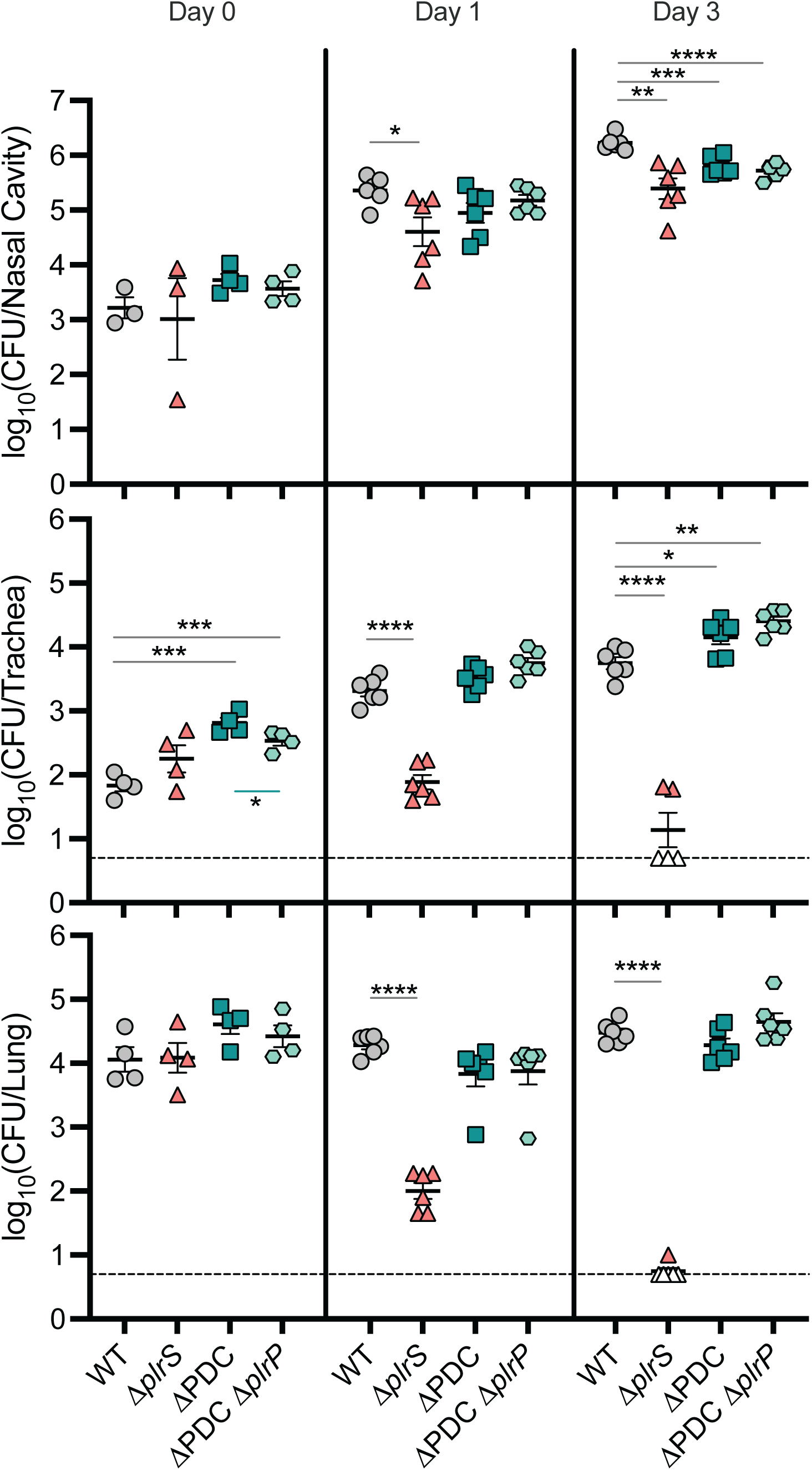
***B. bronchiseptica* persistence in the murine respiratory tract does not require the PlrS PDC domain** Bacterial burden over time within the nasal cavity (top), trachea (middle), and right lung lobes (bottom) of mice infected with 7.5×10^4^ CFU of wild-type (WT), Δ*plrS*, ΔPDC, or ΔPDCΔ*plrP* bacteria. n=4 for day 0, and n=6 for days 1 and 3, with each symbol representing a single mouse. Dashed line represents the limit of detection. Empty symbols represent symbols below the limit of detection. Statistical significance, as determined using unpaired Student’s t-test, is indicated as *, p< 0.05; **, p<0.01; ***, p<0.001; ****, p<0.0001.

### Evidence that PlrR must be phosphorylated at a high level *in vivo*

Although the Δ*plrS* and PlrS_H521Q_ strains grow *in vitro*, they are cleared rapidly from the LRT (17), suggesting that PlrR∼P is required for bacterial survival in the LRT. The Δ*plrS* strain producing PlrR with the D52E phosphomimetic substitution (Δ*plrS* PlrR_D52E_), is viable *in vitro* and, unlike the Δ*plrS* strain, the Δ*plrS* PlrR_D52E_ strain adheres to L2 cells at levels similar to wild-type bacteria when incubated in the presence of 5% CO_2_, indicating that PlrS activity is not required for PlrR activity *in vitro* if PlrR contains the D52E phosphomimetic substitution (18). These data suggest that PlrR is phosphorylated to some extent *in vitro*, and this level of phosphorylation is required for the increased adherence that occurs in the presence of CO_2_ (18). In the murine model, the Δ*plrS* PlrR_D52E_ strain was defective for persistence in the LRT relative to wild-type bacteria (Fig. 6), although not as defective as the Δ*plrS* mutant, suggesting that PlrR_D52E_ is not as active as PlrR∼P and that high levels of PlrR∼P are required *in vivo*.

**Figure 6.**
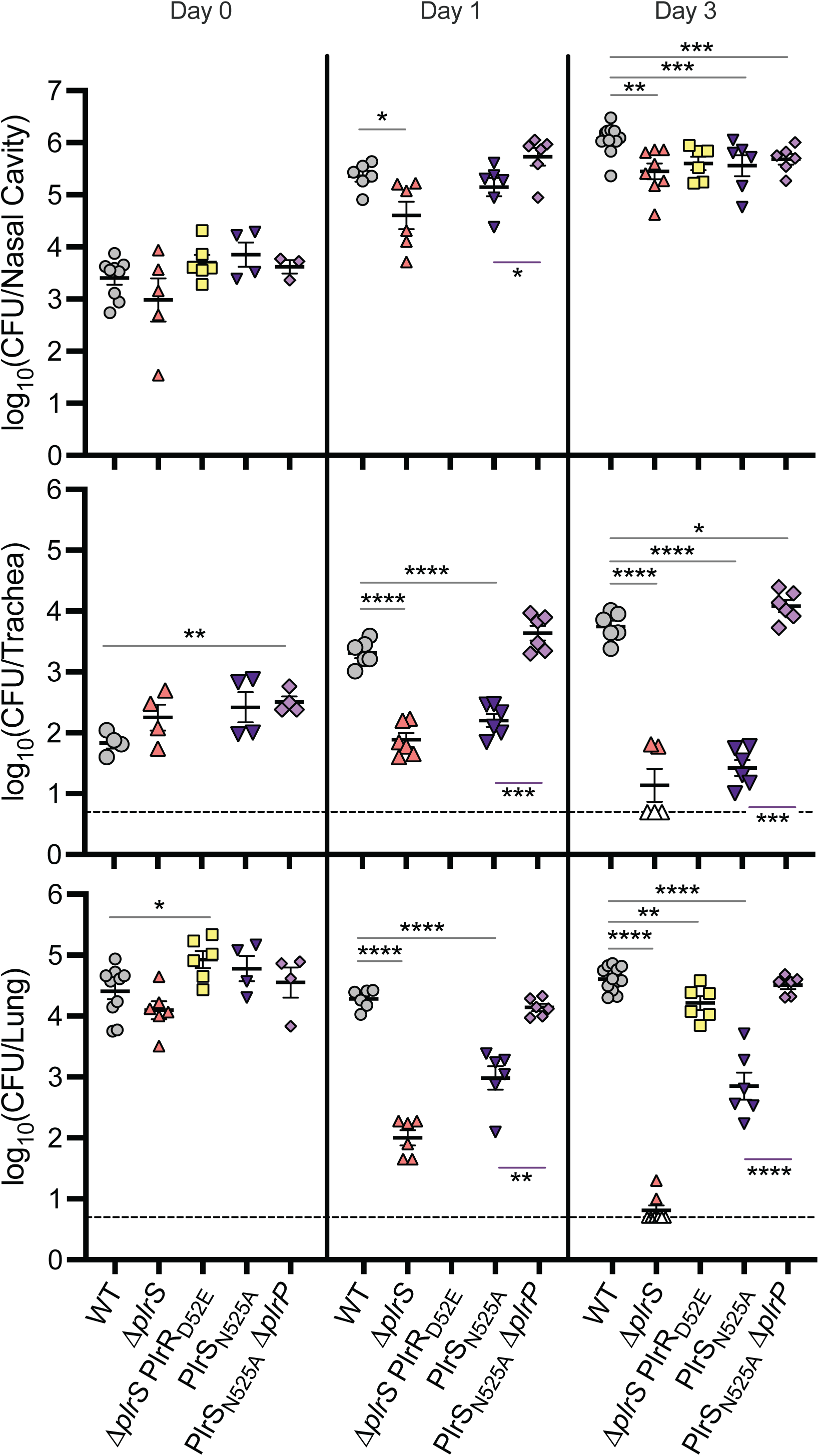
PlrR_D52E_ is not as active as PlrR∼P and PlrP affects PlrS activity in the murine respiratory tract. Bacterial burden over time within the nasal cavity (top), trachea (middle), and right lung lobes (bottom) of mice infected with 7.5×10^4^ CFU of wild-type (WT), Δ*plrS*, PlrS_N525A_, PlrS_N525A_Δ*plrP*, or Δ*plrS* PlrR_D52E_ bacteria. These data are compiled from two independent experiments; each performed in biological duplicate. For experiment 1, which includes WT, Δ*plrS*, PlrS_N525A_, and PlrS_N525A_Δ*plrP* strains, n=4 for day 0, and n=6 for days 1 and 3. For experiment 2, n=6 for days 0 and 3 for the WT and Δ*plrS* PlrR_D52E_ strains, and n=2 for days 0 and 3 for the Δ*plrS* strain. Each symbol represents a single mouse. Dashed line represents the limit of detection. Empty symbols represent symbols below the limit of detection. Statistical significance, as determined individually for each data set using unpaired Student’s t-test, is indicated as *, p< 0.05; **, p<0.01; ***, p<0.001; ****, p<0.0001.

### Evidence that PlrP affects PlrS activity *in vivo*

Because we could not delete *plrP* in wild-type bacteria, we could not determine directly if *plrP* is required during respiratory infection. The PlrS_N525A_ strain is defective for persistence in the LRT, but not as defective as the Δ*plrS* strain ((21), Fig. 6). Because PlrS_N525A_ is defective for phosphatase activity, we previously concluded that both PlrS kinase and phosphatase activity must be required in the LRT. However, the PlrS_N525A_ Δ*plrP* double mutant was recovered from the LRT at levels similar to the wild-type strain at both day 1 and day 3 post-inoculation (Fig. 6). The most likely explanation for these data is that PlrS_N525A_ is defective for both kinase and phosphatase activity, and that the absence of PlrP causes PlrS_N525A_ to have increased kinase activity – further supporting the hypothesis that PlrR∼P levels must be higher in the LRT than when the bacteria are growing in SS medium *in vitro*. Most importantly, these data provide evidence that PlrP affects PlrS activity *in vivo*.

## Discussion

Our data strongly suggest that PlrP is a third component of the PlrSR (NtrYX) subfamily of TCSs and they support the model shown in Fig. 7 and further explored in Supplemental Fig. 3 and 4. According to this model, when *B. bronchiseptica* is grown under standard laboratory conditions *in vitro*, PlrS functions as a weak kinase (or kinase and phosphatase activities are balanced) such that PlrR∼P levels are low. PlrP, likely by interacting with the PlrS PDC domain, prevents PlrS from acting as a strong phosphatase against PlrR∼P. Our data indicate that PlrR∼P is essential *in vitro,* and that in the absence of PlrS kinase activity, PlrR can obtain a phosphoryl group from another molecule. According to our model, a high level of PlrR∼P, which is dependent on PlrS kinase activity, is required in the LRT during infection. PlrP likely also affects PlrS activity *in vivo*, although its exact role *in vivo* is unclear.

**Figure 7.**
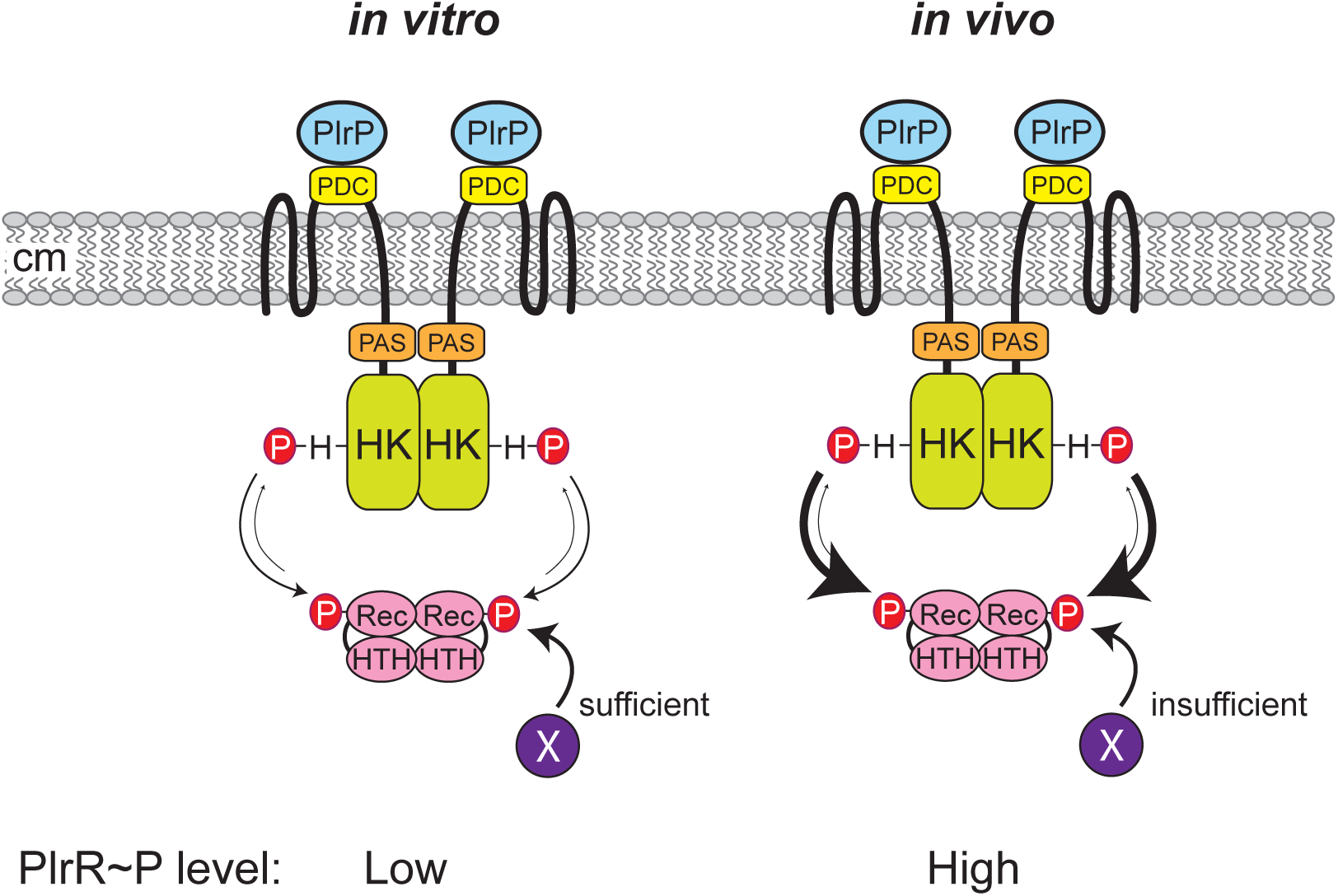
**Model for PlrPSR activity *in vitro* and *in vivo*** According to this model, PlrS functions as a weak kinase when *B. bronchiseptica* is growing in SS medium *in vitro*, and PlrP functions to prevent PlrS from acting as a phosphatase. Another phosphodonor (X) can support PlrR phosphorylation if PlrS is mutated. When *B. bronchiseptica* is growing the LRT (*in vivo*), PlrS has strong kinase activity towards PlrR and PlrR∼P levels are high. Although PlrP affects PlrS activity in the LRT, its role is unknown.

Attempts to delete *plrR* have been unsuccessful (17, 18), and it was unknown whether PlrR (i.e., non-phosphorylated PlrR) or PlrR∼P is essential *in vitro*. Because strains in which PlrS cannot autophosphorylate, such as the Δ*plrS* and PlrS_H521Q_ strains, are viable, either PlrR is essential, or PlrR must be able to obtain a phosphoryl group from another molecule *in vitro*. Our new data showing that *plrP* can be deleted in the PlrS_N525A_ strain, which is defective for phosphatase activity, but not in the wild-type strain, suggests that PlrP prevents strong phosphatase activity by PlrS, which is lethal *in vitro*, presumably due to the dephosphorylation of PlrR. Therefore, the PlrS_H521Q_ strain must also be defective for phosphatase activity. The fact that *plrP* can also be deleted in the PlrR_D52E_ strain, in which the mimicked phosphorylation state of PlrR is independent of PlrS, indicates that the essentiality of *plrP* is dependent on the phosphorylation state (or mimicked phosphorylation state) of PlrR. Together, these data indicate that PlrR∼P is essential *in vitro*, and that PlrR must be able to obtain a phosphoryl group from another molecule *in vitro*.

How does PlrP prevent strong phosphatase activity by PlrS? Because PlrP is predicted to contain a signal sequence for export to the periplasm, we hypothesized that PlrP affects PlrS activity via its periplasmic PDC domain. Since DUF4390 composes the majority of PlrP (161 out of the 168 amino acids of the mature peptide), it is likely that this domain is involved in the interaction. The fact that *plrP* can be deleted in the ΔPDC strain suggests that the essentiality of PlrP depends on the PlrS PDC domain, and that the PDC domain controls PlrS phosphatase activity *in vitro*. While our genetic analyses suggest that PlrP interacts with the PlrS PDC domain in the periplasm to prevent phosphatase activity, further biochemical analyses are required to determine colocalization and direct versus indirect interactions. These analyses would also elucidate the function of DUF4390, a widely distributed but as-yet undefined domain.

Based on previous studies, replacement of the Asp that is the primary site of phosphorylation in a response regulator protein with Glu, a phosphomimetic, results in a response regulator that is at least partially active (44). We showed previously that the *in vitro* adherence defect of a Δ*plrS* mutant is complemented by PlrR_D52E_, suggesting that PlrR_D52E_ is active *in vitro* (18). Although the Δ*plrS* PlrR_D52E_ strain was not as defective as the Δ*plrS* strain in the LRT, it did not persist as well as the wild-type strain, suggesting that PlrR_D52E_ is not as active as PlrR∼P and that high levels of PlrP∼P are required for persistence in the LRT. Because the PlrS_H521Q_ mutant is viable *in vitro* but not *in vivo*, the predicted alternate phosphodonor for PlrR must either not be present when the bacteria are in the LRT, or, more likely, it is unable to produce sufficient levels of PlrR∼P for *in vivo* survival. Together, these data suggest that while relatively low levels of PlrR∼P are sufficient *in vitro*, high levels of PlrR∼P are required *in vivo*.

Determining the exact role of PlrP *in vivo* is a challenge, since we cannot delete *plrP* in the wild-type strain *in vitro*. Our results suggest that neither the PlrS PDC domain nor PlrP in the ΔPDC mutant are required for *B. bronchiseptica* persistence in the LRT (Fig. 6), but in the PlrS_N525A_ mutant, PlrP must affect PlrS activity *in vivo* (Fig. 6). While the N525A substitution results in PlrS that cannot dephosphorylate PlrR (as predicted), PlrS_N525A_ is also slightly defective for kinase activity (21). Therefore, the persistence defect of the PlrS_N525A_ mutant *in vivo* could be due to reduced kinase activity. In which case, PlrS_N525A_ in the absence of PlrP has kinase activity sufficient for *B. bronchiseptica* persistence in the LRT. These results are puzzling yet informative, as we continue to demonstrate that PlrSR activity significantly differs between *B. bronchispetica in vitro* and *in vivo* growth conditions.

*plrP* is not unique to *Bordetella bronchiseptica.* Not only are homologs of *plrP* found in the genomes of many β- and γ-proteobacteria, they are also almost exclusively found 5’ to *plrSR* homologs. The conserved structure of this gene cluster indicates a conserved functional link between these genes. Therefore, it is likely that other PlrP homologs modulate the kinase and/or phosphatase activity of their cognate PlrS homologs.

PlrP is not the first protein proposed to contribute to the function of an NtrYX-family TCS. In the ⍺-proteobacterium *Caulobacter vibrioides,* NtrZ, a putative periplasmic protein, was found to regulate the levels of phosphorylated NtrX through NtrY, presumably by reducing NtrY phosphatase activity (29). Unlike *plrP, ntrZ* is not encoded adjacent to *ntrYX* and is not predicted to encode a DUF4390-containing protein. Additionally, NtrZ has only 15.1% sequence identity with PlrP. However, COBALT comparison of PlrP and NtrZ indicated conservation between the two proteins that includes all the amino acids encoded by NtrZ (45). This result opens the possibility that utilization of a periplasmic regulatory protein is common in NtrYX-family TCSs, even outside of proteobacteria.

Our results indicate that *plrP* is essential in wild-type *B. bronchiseptica in vitro* but possibly not *in vivo*. Given the essentiality of PlrSR during infection and the proposed interaction between PlrP and PlrSR, this finding was unexpected. However, NtrYX- family TCSs are not exclusively linked to pathogenesis. In many bacteria, NtrYX-family TCSs regulate critical cellular functions, including nitrogen fixation and response to oxygen tension (22, 23, 46). PlrP is likely important for responding to a signal that *B. bronchiseptica* does not encounter within a mammalian host but may encounter when living in the environment. Similarly, *rsmB*, which was very frequently found 5’ of *plrP* and *plrSR,* could contribute to survival outside a host. Alternatively, since *rsmB* expression remained below the detection limit under all the conditions we tested*, rsmB* may have been conserved due to the promoter in its 3’ end.

We began this analysis of the gene cluster surrounding *plrSR* in part to parse what contributed to the difficulty of performing genetic manipulations within this region. Through RNAseq analysis and promoter-*gfp* fusion assays, we have determined that at least under laboratory conditions, this cluster does not function as an operon. Instead, there are two promoters for this region, beginning 5’ of *plrP* (P*_plrP_*) and 5’ of *plrR* (P*_plrR_*).

The localization of these two promoters allows for differential regulation of *plrS* and *plrR*. In fact, under laboratory conditions, P*_plrR_* was a stronger promoter than P*_plrP_* (Fig. 3). The decoupling of *plrS* and *plrR* expression indicates that different stoichiometries of PlrS and PlrR are required under different conditions. With these promoters identified, it will be possible to generate mutations that maintain regulation through this region, allowing deeper analysis of this unusual but clinically important two-component system.

## Materials & Methods

### Ethics statement

This study was carried out in strict accordance with the *Guide for the Care and Use of Laboratory Animals* of the National Institute of Health. Our protocol was approved by the University of North Carolina Institutional Animal Care and Use Committee (ID: 22–140). All animals were anesthetized for inoculations, monitored daily, and properly euthanized. All efforts were made to minimize suffering.

### Bacterial culturing

*Bordetella bronchiseptica* strains were grown on Bordet-Gengou (BG) agar plates (BD Biosciences) supplemented with 6% defibrinated sheep blood (Hemostat) at 37°C for 2-3 days. *B. bronchiseptica* strains were grown in Stainer-Scholte (SS) broth supplemented with SS supplement ((47), updated in (48)) at 37°C on a rotating wheel to increase aeration overnight or until the desired density was reached. *E. coli* strains were grown in Lysogeny Broth (LB) at 37°C on a rotating wheel overnight or on LB agar plates at 37°C for 1-2 days. As needed, media was supplemented with streptomycin (Sm, 20 μg/mL), gentamicin (Gm, 30 μg/mL), kanamycin (Km, 50 μg/mL), diaminopimelic acid (DAP, 300 μg/mL), ampicillin (Ap, 100 μg/mL), or sucrose (15% w/v). All cultures were started from individual colonies from a clonal population when possible.

### Construction of plasmids and strains

The strains and the plasmids that were used in this study can be found in Supplemental Table 1. In-frame deletions were constructed via allelic exchange using derivatives of the pEG7S plasmid (49). Transcriptional reporter strains were constructed via transposase-mediated insertion at the *att*Tn*7* site using derivatives of the pUC*gfp*MAB vector (50). Plasmids were constructed and propagated within the DH5a *E. coli* strain. The RHO3 *E. coli* strain was used for introducing plasmids into *B. bronchiseptica* through conjugation. Plasmids were confirmed using sequencing and all mutations introduced in *B. bronchiseptica* strains were confirmed by PCR.

### Bioinformatic analysis of the *plrSR* gene cluster

The gene neighborhoods of the genes encoding NrYX-family TCSs outlined in Table 1 were aligned using CAGECAT clinker (39). Unbiased examination of the gene cluster was performed using *fast.genomics* (40). This database uses *Bordetella pertussis* strain 18323 as the representative *Bordetella* strain. We used the “gene neighborhood tool” to identify the closest homologs to *plrS* (BN118_RS17975; 200 top hits, 9kb neighborhood). This dataset was manually examined for neighborhood structure similarities, with samples being removed if they lacked complete sequencing across the 9kb region. We used the “compare gene presence/absence” tool to examine the co-occurrence of *plrS*, *plrP* (BN118_RS17980), and *rsmB* (BN118_RS17975). For *plrs/plrP*, the table of the best homologs of both genes was used for further analysis. The gene neighborhoods of the good *plrS* homologs were manually examined for the presence of an adjacent gene encoding a DUF4390-containing protein. AnnoTree (AnnoTree v2.0.0; GTDB Bacteria Release R214) was used to visualize the distribution of *plrP* (Pfam ID: PF14334) and *rsmB* (Pfam ID: PF01189) distribution across proteobacteria (41).

**Table 1.** Genes compared using CAGECAT clinker.

| Species | Accension or Assembly number | Gene IDs |
| --- | --- | --- |
| *Azorhizobium caulinodans* | NC_009937.1 | AZC_3081-3088 |
| *Azospirillum brasilense* | ASM782742v1 | OH82_RS30760-RS30810 |
| *Bordetella bronchiseptica* | NC_002927.3 | BB0262-0267 |
| *Bradyrhizobium diazoefficiens* | NZ_CP011360.1 | AAV28_RS19055-RS19085 |
| *Brucella abortus* | NC_006932.1 | BruAb1_1120-1125 |
| *Caulobacter vibrioides* | NC_002696.2 | CC1739-CC1744 |
| *Cereibacter azotoformans* | ASM2139131v1 | LV780_RS06670-RS06710 |
| *Herbaspirillum seropedicae* | NZ_CP011930 | ACP92_RS00340-RS00355 |
| *Neisseria gonorrhoeae* | NZ_AP023069 | HT085_RS10140-RS10180 |
| *Paracoccus denitrificans* | ASM20389v1 | Pden_4123-4131 |
| *Rhodobacter sphaeroides* | ASM1290v2 | RSP_2836-2844 |
| *Sinorhizobium meliloti* | NZ_CP146208 | V7S97_RS16985-RS17025 |

### RNA sequencing analysis

The RNA sequencing data used in this study has been previously published (42); the raw sequencing files can be retrieved from the GEO repository (GSE268598). Read counts were determined using the igvtools Count command on BAM alignment files within the Integrative Genomic Viewer (51). Counts were mapped to the RB50 genome (RefSeq NC_002927.3).

### *in vitro* promoter activity analysis

The transcriptional reporter strains were grown for 16-18 hours in standard SS medium or SS medium containing 40 mM MgSO_4_ at 37°C. 200 μL of each culture was transferred to a black, clear bottom 96-well plate (ThermoScientific catalog no. 165305), and the OD_600_ and GFP fluorescence (excitation: 485nm, emission: 535nm) were measured on a BioTek Synergy H1 Hybrid Reader plate reader. The relative fluorescence intensity (RFI) was calculated by normalizing GFP fluorescence to OD_600_ and subtracting the empty vector control values. The experiments were performed in biological triplicate.

### Bacterial infection of the mouse respiratory tract

Six-week-old female BALB/c mice from Charles River Laboratories (catalog no. BALB/cAnNCrl) were inoculated intranasally with 7.5 x 10^4^ CFU *B. bronchiseptica* in 50 μL of DPBS. At 0, 1, or 3 days post-infection, the right lung lobes, the trachea, and nasal cavity tissues were harvested from each mouse. The tissues were homogenized in DPBS using a mini-beadbeater with 0.1 mm zirconia beads (Biospec catalog no. 11079110zx). The number of CFU was determined by plating dilutions of tissue homogenates on BG Sm blood agar and enumerating the number of colonies per tissue after at least 48 hours of growth at 37°C.

## Acknowledgements

This work was supported by funding from the NIH (R01 AI153160 to PAC and R01 AI129541 and R21 AI177818 to PAC and SMJ).

